# Altered kinship vocal dynamics in marmosets with valproic acid–induced model of autism

**DOI:** 10.1101/2025.01.30.635822

**Authors:** Koki Mimura, Keiko Nakagaki, Hirofumi Morishita, Noritaka Ichinohe

## Abstract

Autism spectrum disorder (ASD) is characterised by social communication impairments and repetitive behaviours. Language deficits, including echolalia and restricted vocabulary, heighten caregiver stress and negatively affect the family’s quality of life. Although animal models have advanced the understanding of individual ASD traits, their influence on kinship dynamics remains underexplored. To address this issue, we developed a clinically relevant ASD model in common marmosets by prenatal exposure to valproic acid (VPA) to produce ASD-like pups alongside their typically developing parents. We analysed 28,418 kinship calls from nine VPA-exposed and seven unexposed (UE) pups, along with their parents. Kinship vocalisations in VPA families exhibited significant alterations, including increased isolation calls, decreased affiliative calls, disruption of structured repetition patterns, and reduced developmental maturations. These deviations intensified after weaning, suggesting a link between social communicative stressors and altered family dynamics. Parental weight loss was correlated with kinship vocal deviations, potentially reflecting increased caregiver stress. This observation aligns with clinical reports of heightened stress in families raising children with ASD. VPA pups also displayed premature locomotion independence, indicating broader social and communication disruptions. These findings suggest that VPA marmosets are valuable models for investigating ASD-like traits in individuals and kinship-level dynamics. Kinship vocalisations provide critical insights into the interplay between communication impairments and caregiver stress, offering a promising avenue for developing non-invasive biomarkers for ASD-related challenges.

## Introduction

Autism spectrum disorder (ASD) is a neurodevelopmental condition characterised by difficulties in social communication and restricted and repetitive behaviours.(1) These symptoms typically emerge in early childhood and intensify as social demands increase.(2) Impaired language development in patients with ASD shows characteristic alterations, including echolalia and atypical vocal patterns, refracting broader impairments in communicative flexibility.(3)(4) Such communication deficits not only impact the quality of life of individuals with ASD but also impose considerable stress on caregivers, exacerbating family dynamics.(5)(6)(7) Compared with families of children with other developmental disorders, families with ASD face heightened kinship stress, particularly during critical early developmental social milestones such as weaning and preschool entry.(8)

Despite advances in neuroscience research using animal models of ASD, our understanding of kinship dynamics remains limited. Existing animal models, predominantly rodent models, have been instrumental in elucidating the neural mechanisms underlying ASD.(9) However, rodents lack the social complexity and cooperative kinship structures that characterise human families. To bridge this gap, the common marmoset (*Callithrix jacchus*), a small nonhuman primate, offers distinct advantages. Marmosets possess a kinship system that closely parallels that of humans and is characterised by paternal and elder involvement in childcare, altruistic behaviours, and rich vocal communication. During their early development, they acquire complex social skills such as third-party reciprocity and fairness within the family environment.(10)(11) These unique traits make marmosets a valuable model for investigating the developmental dynamics of kinship systems.

Building on these strengths, we developed a marmoset model of ASD using prenatal exposure to valproic acid (VPA), an antiepileptic drug clinically linked to ASD-like symptoms when exposure occurs during the foetal neural tube development stage.(12) In rodents, prenatal VPA exposure is a well-established model of ASD that results in altered neural development and social impairments.(9) In marmosets, VPA exposure induces ASD-like phenotypes,(13) including atypical neural development,(14)(15)(16) social impairments,(17)(18)(19) and heightened daily stress.(20) The VPA-exposed marmoset model incorporates typically developing parents and ASD-like pups, thus providing a unique framework for investigating how ASD traits affect kinship dynamics over time.

ASD symptoms often emerge during key developmental milestones in marmosets, such as weaning, which occurs between postnatal months 2 and 3 (PM 2–3).(21) This period offers a valuable opportunity to examine behavioural phenotypes and kinship interactions that may reflect ASD-like traits. We hypothesised that kinship behaviours during weaning could serve as sensitive, non-invasive indicators of stress and social communication impairment in marmoset families with VPA-exposed pups (VPA families).

A unique feature of marmosets is their rich vocal repertoire, which earns them the nickname-song monkeys. They produce more than 10 distinct call types,(22) including the isolation-indicating *phee* call(23) and the socially engaging *trill* call.(24) These vocalisations provide a window into social and emotional dynamics, making them particularly suited for studying the interactions between the pup models of ASD and typically developing family members. Existing approaches to vocal communication analysis in animal models often require expensive methods to identify individual callers, limiting their scalability and ecological validity.(23)(25)(26)(27) To address these challenges, drawing on insights from natural language analysis in ASD,(28)(29) we focused on call frequency and sequence patterns, instead of individual identification, to capture the overarching social dynamics within the kinship system.

In this study, we aimed to explore how VPA exposure affects kinship behaviour and stress, particularly during the weaning period. By recording and analysing locomotion, vocalisations, and body weights across marmoset families from PM 1 to 5.5, we sought to uncover atypical patterns associated with ASD traits (Figure 1A). Our approach not only advances the understanding of ASD-related family dynamics but also holds potential for clinical applications, offering insights into stress and communication impairments in human families affected by ASD.

**Fig. 1.**
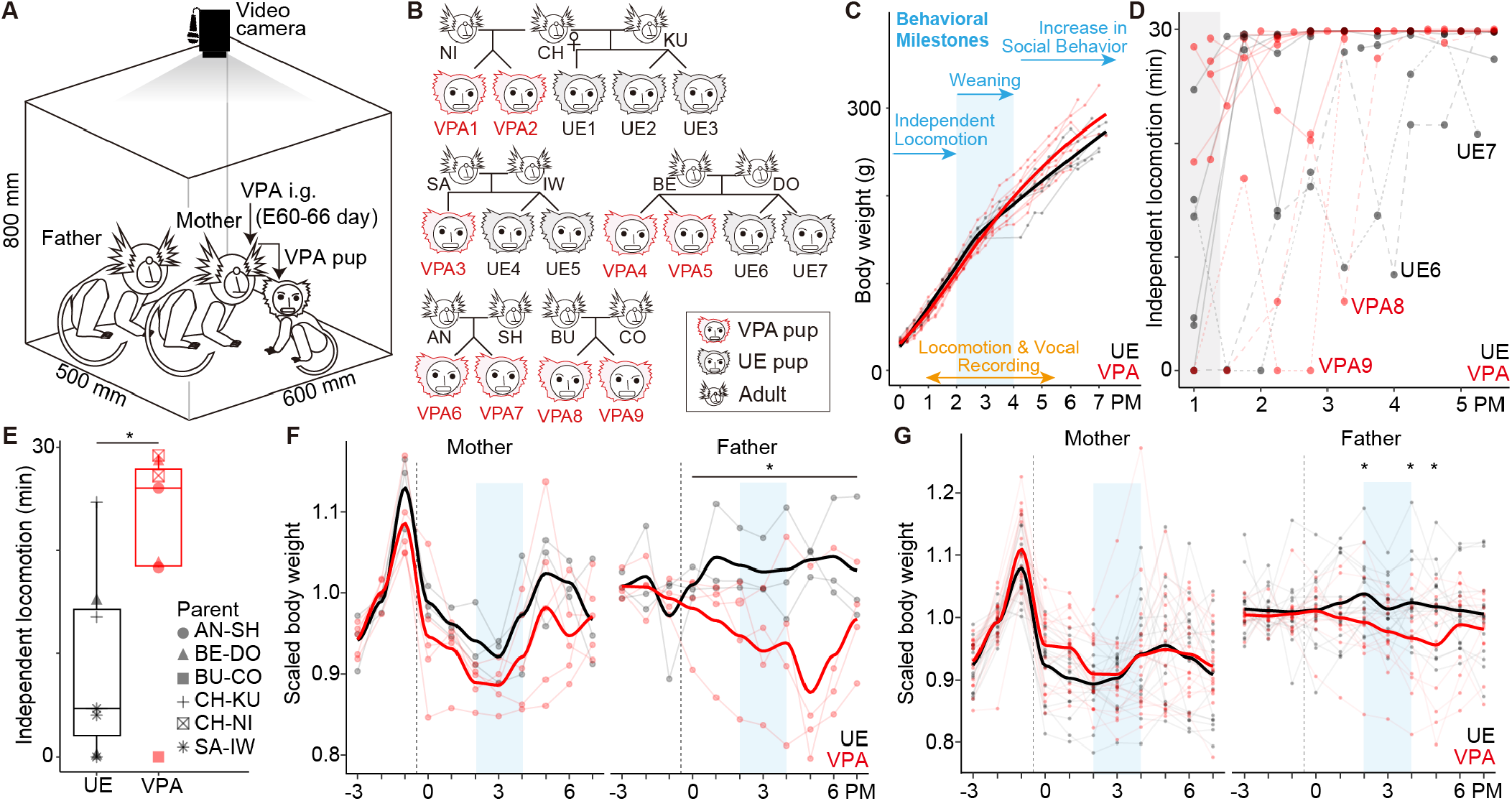
Characteristics of VPA families in terms of behaviour and parental body weight. **(A)** Pups born to mothers that received intragastric (i.g.) administration of VPA during pregnancy are referred to as VPA pups. Kinship behaviours, including locomotion and vocalisations, are recorded by transferring the father, mother, and either a VPA or UE pup in their home cage to a soundproof room. **(B)** Behaviour data included 16 pups (VPA = 9; UE = 7) from five families. The inverted Y-branch represents twin pairs. **(C)** Body weights of pups, overlaid with typical behavioural milestones (top, blue) and the recording period (PM 1–5.5, bottom, orange). **(D)** Scatter plot showing the time pups spent independently during 30-min recording sessions. Two twin pairs, VPA8/VPA9 and UE7/UE6, are highlighted with distinct line types and labelled. **(E)** Boxplots of independent activity duration during the first recording session around PM1 (grey squares in **D**). Markers are differentiated by parental pairs. VPA families exhibit longer independent activity durations than UE families (p < 0.05, Brunner–Munzel test). **(F)** Scatter plots with loess regression lines depicting scaled body weight changes for the fathers and mothers of recorded pups. Body weights are scaled relative to the average during PM −3 to −1. Paternal body weight significantly decreased during the caregiving period for VPA pups (p < 0.05, analysis of covariance). **(G)** Scatter plots with loess regression lines of scaled body weight during the parenting period (VPA = 17; UE = 15) for the same parental pairs. To confirm robustness, the analysis includes data from other caregiving periods experienced by the same parents. Significant decreases in paternal body weight for VPA pups are observed at PM 2, 4, and 5 (p < 0.05, Tukey’s honestly significant difference test). The blue squares in **F** and **G** indicate the weaning period of pups, consistent with those shown in **C**. VPA, valproic acid; UE, unexposed; PM, pups’ age in postnatal months.

## Results

### Early locomotion independence in VPA pups

We investigated the effect of VPA on kinship-level behavioural expression in pups throughout their early developmental stages. Kinship behaviour was recorded using a video camera mounted above the home cage in a soundproof room, allowing one pup and its parents to move freely for 30 min (Figure 1A). This study included nine VPA-exposed pups (seven males and two females) and seven unexposed (UE) pups (five males and two females) as controls (Figure 1B; Table S1). Body weight, monitored as a physical growth indicator, was not significantly different between the groups during PM 0 to 7 (f[1, 13] = 1.482, p = 0.247, repeated-measures analysis of covariance (ANCOVA); Figure 1C). A total of 109 recording sessions (UE = 46; VPA = 63) were conducted biweekly from PM 1 to 5.5 spanning three key behavioural milestones: locomotion independence (∼ PM 2), weaning (PM 2 and 3), and increased social behaviours such as grooming and head-to-head contact between pups and parents (PM 4 onwards; Figure 1C, blue arrows). The first milestone—independent movement away from the parents’ backs—was achieved by PM 3 in most pups, with a few exceptions in both groups (Figure 1D). Interestingly, the effects of VPA exposure were evident as early as the initial recording session around PM 1. VPA pups demonstrated more pronounced independence from their parents than did UE pups (UE = 6, VPA = 7, BM statistic = 2.62, df = 8.46, p = 0.029, Brunner–Munzel test; Figure 1E). In this novel recording context, the reduced tendency of VPA pups to seek a retreat from their parents’ backs or the lack of facilitation by parents suggests potential alterations in their kinship system.

### Paternal body weight loss during the weaning period of VPA pups

To evaluate whether VPA exposure affected not only pup behaviour but also the broader kinship system, parental body weights were continuously monitored. Maternal body weight transitioned from decreasing to increasing around PM 3, corresponding to the weaning period, with no significant effects attributed to VPA exposure (PM 0–7, f[1, 48] = 0.64, p = 0.428, ANCOVA). By contrast, fathers showed a significant reduction in body weight during the caregiving period for VPA pups (PM 0–7, f[1,42] = 16.7, p = 1.96e-4, ANCOVA; Figure 1F). To confirm the robustness of this trend, a systematic analysis was conducted on parental body weight during caregiving, including data from births not associated with the recorded sessions (VPA = 13, UE = 15; Table S2). This analysis replicated this pattern, with significant reductions in paternal body weight observed at PM 2 (estimate = −0.046, adjusted p = 0.048), 4 (estimate = −0.057, adjusted p = 0.022), and 5 (estimate = −0.062, adjusted p = 0.025, Tukey’s honestly significant difference (Tukey-HSD) test; Figure 1G). Unlike maternal weight, which is influenced by factors such as childbirth, lactation, and subsequent pregnancies, paternal weight loss is less likely to be affected by these confounders. These findings suggest that caregiving for VPA pups may require more effort from fathers than caregiving for UE pups. This weight loss trend became apparent after the pups achieved independent locomotion behaviour, suggesting the need to investigate additional metrics to better understand the changes in kinship dynamics during this period.

### The number of kinship calls in VPA families did not decrease with pup development

A total of 28,418 calls were systematically collected from kinship behaviour records, providing a dataset of vocalisations, recognised as a commonly used non-invasive indicator for assessing collective emotional states and enabling longitudinal comparisons. All vocalisation analyses were conducted without identifying the subject that produced the calls. In UE families, the frequency of kinship calls decreased with pup development (n = 46, f[1,33] = 5, p = 0.032, F-test), which is consistent with previous studies reporting a decline in pup call frequency under isolated conditions as development progressed. However, this decline was not observed in VPA families, where the call frequency remained stable over time (n = 63, f[1,45] = 0.03, p = 0.954, F-test; Figure 2A). These results from the simple regression models were supported by linear mixed model (LMM) analysis, which was conducted to examine the model while accounting for the random effects of individual and parental differences. Based on model selection using the Akaike information criterion (AIC), the best model for UE families showed a negative correlation with PM, including the random effects of individual differences (model #13; Tables S3 and S4), with a fixed effect of −81.3. Model 13, the best model, was validated using a likelihood ratio test (LRT) against model 4, which excluded the treatment effect, confirming a significant improvement in the data explanation (deviance residual (DR) = 12.7, df = 1, p = 3.61e-4, LRT). For VPA families, a negative correlation with PM was observed, including the random effects of parental differences (model #14, DR = 9.30, df = 1, p = 0.0229, LRT vs. model #5), but the fixed effect was significantly smaller at −0.186. The lack of a typical developmental decrease in the kinship vocal frequency and pup development in VPA families suggests potential alterations or immaturity in the rules of vocal usage.

**Fig. 2.**
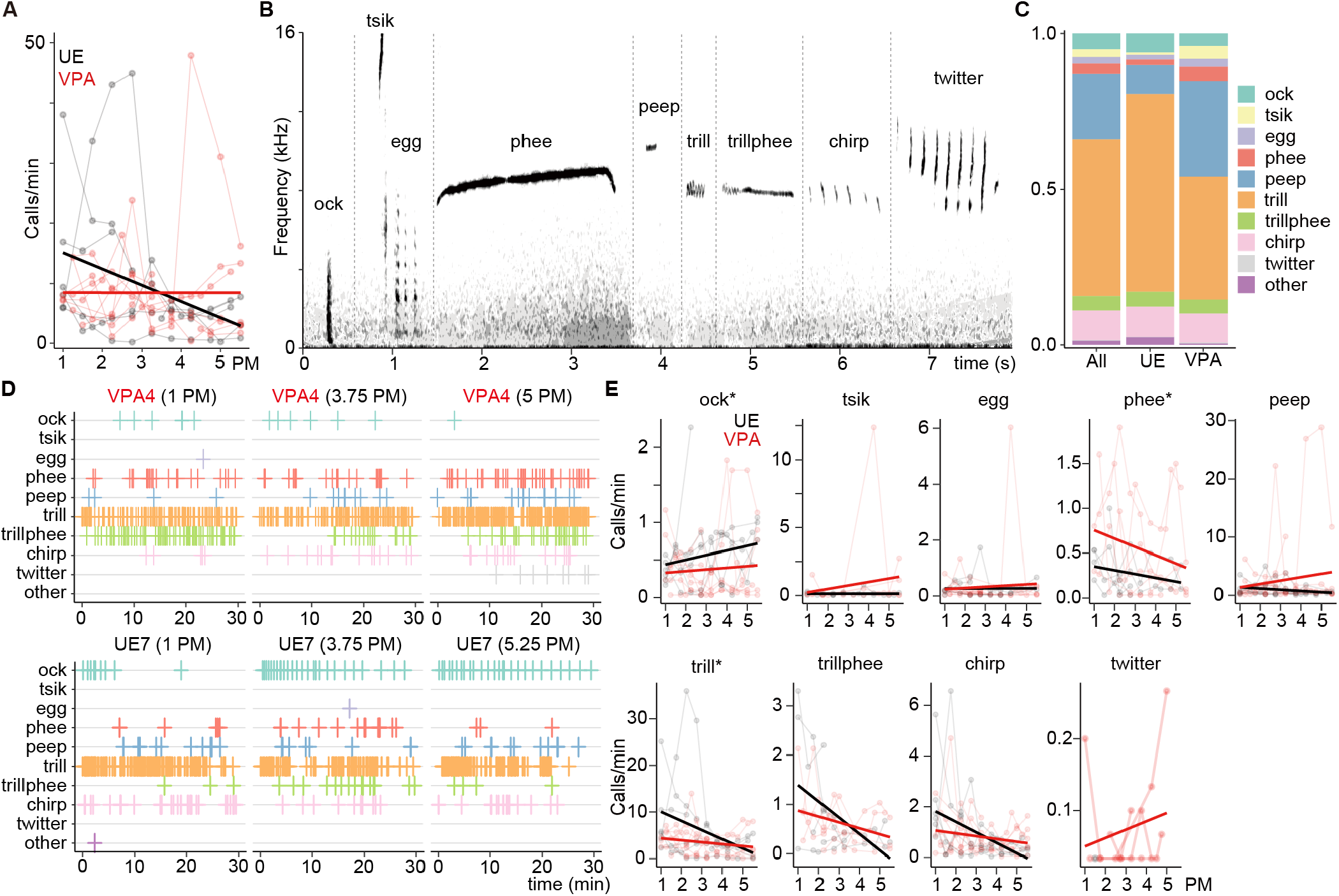
Abnormal kinship vocal call frequencies in VPA families. **(A)** Scatter plot showing the number of kinship vocal calls. Data points are connected by thin lines for each measured family pair. **(B)** Typical spectrogram of the nine major call types. These data are sampled from individual datasets separated by dotted lines and combined for visualisation (see Table S2). **(C)** Call type proportions aggregated across all ages. **(D)** Examples of kinship vocalisations recorded during 30 min of free activity for VPA4 (top) and UE7 (bottom) pups, along with their parents (BE-DO in Figure 1B). The x-axis shows timestamps, and the y-axis indicates call types. Columns represent recordings at PM 1, 3.75, and 5 or 5.25. **(E)** Scatter plots of call frequency. Significant atypical trends are detected in VPA families for three types of calls: ock, phee, and trill (*, p < 0.05, analysis of covariance) (see Table S5). VPA, valproic acid; UE, unexposed; PM, pups’ age in postnatal months.

### Affiliative calls decreased while anxiety-related calls increased in VPA families

To evaluate the social stress conditions in VPA families, we identified diverse call types within kinship vocalisations and focused on the frequency of calls linked to emotional states. Excluding 125 unclassifiable calls, the remaining calls were categorised into nine distinct types: *ock, tsik, egg, phee, peep, trill, trillphee, chirp*, and *twitter* (Figure 2B; Table S5). Of all calls, *trill*—known as a social affiliative call—was the most frequent at 50.3% (n = 14,426), whereas *phee*—known as an anxiety-related call—accounted for only 3% (n = 955). No significant differences were observed in the composition ratios of calls pooled across the entire measurement period between the UE and VPA families (statistic = 0.225, df = 9, p = 1.00, chi-square test; Figure 2C). However, the visualisation of representative examples revealed distinct distributions of call types, with an increase in *phee* observed in VPA families and an increase in *ock* in UE families (Figure 2D). Further statistical comparisons of call frequencies by type highlighted notable differences: in VPA families, the frequency of *trill* significantly decreased (f[1,102] = 5.56, p = 0.020, ANCOVA), whereas that of *phee* significantly increased (f[1,68] = 7.50, p = 0.008, ANCOVA). A significant negative correlation with PM was also observed for *trill* (f[1,102] = 8.64, p = 0.004, ANCOVA). Additionally, notable changes were observed in ambiguous calls such as *ock* and *twitter*. In VPA families, the frequency of *ock* significantly decreased (f[1,98] = 5.71, p = 0.019, ANCOVA) and most of *twitter* (38/39) calls were observed in VPA families (Figure 2E). These changes in call types were supported by the LMM analysis, where significant VPA exposure effects were detected across *trill, phee*, and *ock*. Random effects were also observed for pups in terms of *trill* (model #17 in Table S3 selected by LMM analysis, DR = 17.9, df = 2, p = 1.30e-4, LRT vs. model #13) and for parents in terms of *phee* and *ock* (model #18; phee, DR = 21.5, df = 2, p = 2.16e-5; ock, DR = 7.72, df = 2, p = 2.11e-2, LRT vs. model #14; Tables S3 and S6). The frequency of representative calls, which is known to be associated with emotional states, showed a consistent trend of increased social stress and decreased affiliative mood throughout the measurement period.

### Call repetition frequency of kinship vocalisation was altered in VPA families

Given that impairments in language communication in ASD are characterised by echolalia or unnatural repetition of words, leading to reduced communicative flexibility, we examined the sequential organisation of calls in kinship vocalisations. Specifically, we analysed all pairs of calls with inter-call intervals (ICIs) ≤ 30 s (Figure 3A), categorising them as either repeated or non-repeated and further classified into short ICIs (ICI ≤ 10 s) or long ICIs (10 < ICI ≤ 30 s). Thresholds of 10 s and 30 s corresponded approximately to the 85th and 96th percentiles of the ICI distribution, respectively (Figure 3B). In UE families, the number of short ICI repeated calls decreased with pup development (n = 5539, f[1,43] = 15.8, p = 2.69e-4, F-test), whereas that of long ICI non-repeated calls increased (n = 1289, f[1,43] = 9.75, p = 0.003, F-test) (Figure 3C). These developmental trends were supported by the LMM analysis, with model #10 being the best fit (Table S3). By contrast, no such trends were observed in VPA families (short ICI repeated: n = 7017, f[1,61] = 0.02, p = 0.89, F-test; model 5, DR = −8.40, df = 1, p = 1.00, LRT vs. model #14; long ICI non-repeated: n = 2213, f[1,61] = 0.401, p = 0.53; model #1). These findings suggest a disruption of sequential rules in the kinship vocalisations of VPA families, diverging from the structured developmental patterns observed in UE families (Tables S3 and S7).

**Fig. 3.**
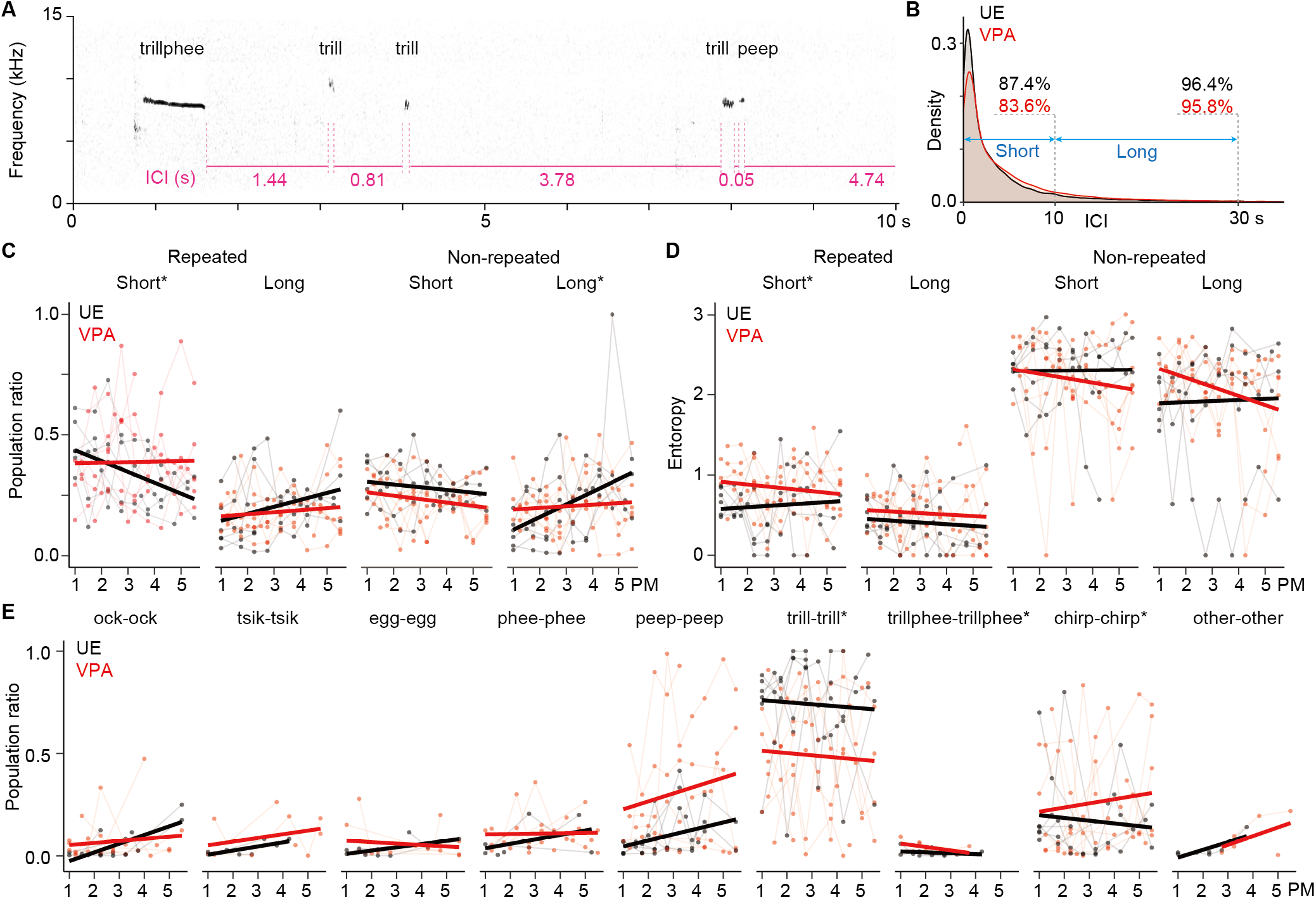
Alteration of call repetition rules in VPA families. **(A)** Example spectrogram of kinship vocalisations from UE2 and its parents at 1.5 PM. The inter-call interval (ICI) is defined as the time between the end of a pre-call and the start of a post-call. **(B)** Density distribution of the ICI. Two calls with ICI ≤ 30 s are classified as related calls and further divided into short (ICI ≤ 10 s) and long (10 < ICI ≤ 30 s) intervals. The percentiles of the ICI for UE and VPA families are indicated above the density plot in black and red, respectively. **(C)** Longitudinal comparison of call repetition rules during development. Related calls are categorised into four types based on the ICI (short or long) and whether they are repetitions of the same call type (repeated) or different types (non-repeated) (see Tables S6 and S7). **(D)** Entropy of 2-call phrases categorised by repetition and ICI. **(E)** Proportional representation of each call type in repeated calls within a short ICI (≤ 10). Significant VPA exposure effects are observed for three repeated call types based on LMM analysis: *trill*-*trill* (UE > VPA), *trillphee*-*trillphee* (UE < VPA), and *chirp*-*chirp* (UE < VPA) (see Table S7). VPA, valproic acid; UE, unexposed; PM, pups’ age in postnatal months.

### Disruption of structured repetition patterns in short ICI calls of VPA families

To characterise the compositional changes in repeated calls, we analysed the distribution patterns of 2-call phrases, focusing specifically on short ICI repeats. Entropy analysis (average information, H) of the four categories of 2-call phrases revealed significantly higher diversity in VPA families, reflecting a departure from the organised vocal structure observed in UE families (f[1,102] = 10.18, p = 0.002, ANCOVA; model #6 in Table S3; Figure 3D). A detailed analysis of the 10 call types (9 categories plus ‘other calls’) revealed specific differences in repetition patterns. In UE families, *trill*-*trill* accounted for approximately 75% of the short ICI repeats, whereas, in VPA families, this proportion significantly decreased to approximately 50% (f[1,95] = 22.2, p = 8.43e-6, ANCOVA; model #9). Conversely, VPA families exhibited significant increases in *trillphee*-*trillphee* (f[1,16] = 5.41, p = 0.034, ANCOVA; model #15) and *chirp*-*chirp* (f[1,76] = 12.8, p = 6.13e-4, ANCOVA; model #6) repeats. Although the frequency of *peep*-*peep* was higher in UE families than in VPA families (f[1,78] = 12.8, p = 6.13e-4, ANCOVA), the random effects of parental differences absorbed this variation, resulting in no significant group differences in the LMM analysis (model #5) (Figure 3E; Table S8). These findings demonstrate that VPA families deviate from socially meaningful and consistent repetition patterns, such as *trill*-*trill*, commonly observed in UE families. Instead, VPA families exhibited an increase in non-affiliative repetitions. This breakdown of structured vocal patterns parallels ASD-like echolalia, which is characterised by atypical repetitive vocal usage in social communication. Collectively, these results highlight how VPA exposure disrupts the organisation of socially relevant vocal sequences, leading to more variable and less structured vocal interactions.

### Reduced developmental changes in multi-call phrase frequency distributions in VPA families

VPA families exhibited developmental delays or stagnation in kinship vocalisations, with significantly smaller changes in the frequency of single calls and 2-call phrases with specific repetition rules. Within-subject longitudinal comparisons were performed to validate this pattern. Data were divided into three developmental stages with balanced recordings from VPA families: stage 1 (PM 1–2.5, UE = 18, VPA = 21), stage 2 (PM 2.5–3.5, UE = 15, VPA = 21), and stage 3 (PM 3.5–5.5, UE = 13, VPA = 21). Jensen–Shannon divergence (JSD) was calculated to evaluate the changes in the frequency distributions of single calls and multi-call phrases (up to six consecutive calls, ICI ≤ 30). Within-subject comparisons showed a significantly lower JSD in VPA families than in UE families for 4-to 6-call phrases between stages 2 and 3 (p < 0.05, Brunner–Munzel test). No significant differences were observed between stages 1 and 2 (Figure 4A). These findings indicate that focusing on multi-call phrase frequency distributions highlights developmental stagnation in kinship vocal usage in VPA families, which is evident in the form of reduced changes within individuals, particularly during later weaning stages.

**Fig. 4.**
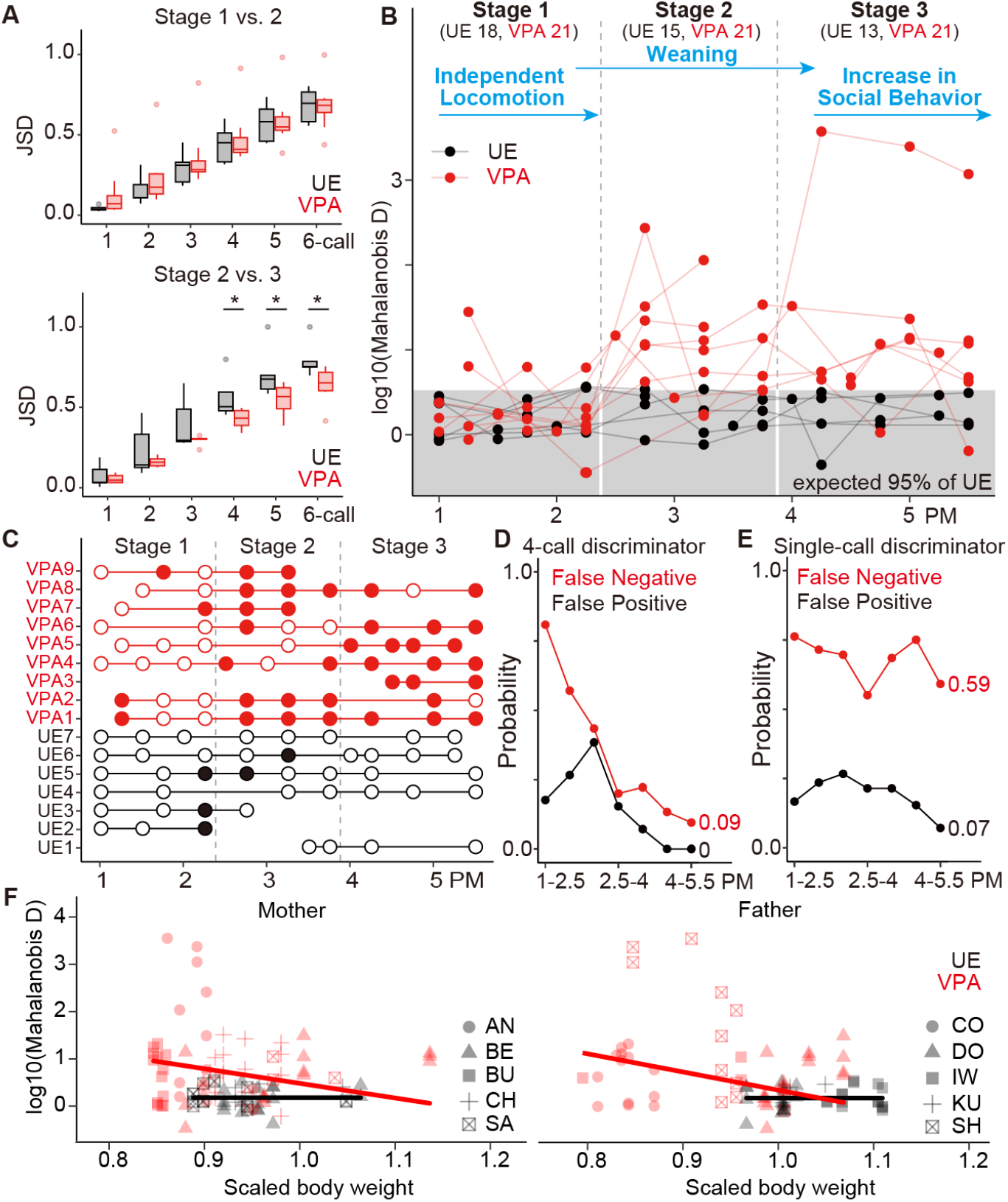
Discrimination of VPA families based on kinship vocal patterns. **(A)** Boxplots showing the expected changes in entropy (JSD) with the development of the related call sequence frequency. The sequences were classified as 1-to 6-call phrases based on the number of consecutive calls (ICI ≤ 10). Comparisons are made between developmental stage 1 (PM 1–2.5) and stage 2 (PM 2.5–4; top) and between stages 2 and 3 (PM 4–5.5; bottom). Significant differences are observed between stages 2 and 3 for 4-call (BM statistic = −3.12, df = 8.94, p = 0.012), 5-call (BM statistic = −3.12, df = 8.94, p = 0.012), and 6-call phrases (BM statistic = −4.19, f = 8.93, p = 2.3e-3, Brunner-Munzel test). **(B)** Mahalanobis distance from UE families for the frequency distribution of 4-call phrases is calculated based on the top 1–5 PC scores across the three developmental stages. The number of data points included in each developmental stage is shown at the top. These stages correspond to typical behavioural milestones in marmosets (blue arrows; see Table S8). **(C)** Results of the discriminant analysis, showing whether data points deviate from the expected 95% distribution range of UE families (grey box in **B**). Outlier data points are filled in, whereas non-deviating points are displayed as a hollow. **(D and E)** Validation of discriminant models to determine UE or VPA. False negative rates (red) and false positive rates (black) are calculated from **C (D)** or from the frequency of single calls **(E). (F)** Scatter plots with regression lines of the best model for the correlation between the scaled parental body weight (Fig. 1F) and log10 of Mahalanobis distance in **B** (see Table S11).VPA, valproic acid; UE, unexposed; ICI, inter-call interval; PC, principal component; PM, pups’ age in postnatal months; JSD, Jensen–Shannon divergence.

### Developmental deviations in 4-call phrasing in VPA families

The finding that multi-call phrase frequencies prominently reflect the characteristics of VPA families opens up new avenues for their use as biomarkers to distinguish families with ASD-like pups. To test this potential, we performed a discriminant analysis focusing on the frequency of 4-call phrases. When aggregating representative 4-call phrases, 4-*trill* phrases were the most frequent across all developmental stages in UE families. However, in VPA families, 4-*chirp* phrases and 4-*peep* phrases consistently showed high frequencies, whereas these phrases tended to be less frequent in UE families during stages 2 and 3 (Table S9). From the dataset, 1713 unique 4-call phrases were identified, and their count data were reduced to five dimensions (explaining 98.45% of the variance; Table S10) using principal component analysis (PCA). We calculated the Mahalanobis distances from the UE mean vector to the individual data across the three developmental stages. This enabled the assessment of the deviation of each data point (Figure 4B). Deviations were defined as exceeding the 95th percentile of the UE distribution and were observed in 63.5% (40/63) of VPA families, with a tendency to increase with pup development (Figure 4C). The false negative rate, indicating the proportion of VPA families indistinguishable from UE families, was the highest during stage 1 at 81% (17/21). However, this value steadily declined as the pups developed, reaching 9.5% (2/21) in stage 3 (Figure 4D, indicated by red). The false positive rate, which indicates the proportion of UE families classified as not UE families, progressively declined and reached zero in stage 3 (Figure 3D, indicated by black). For comparison, a similar discriminant analysis was conducted using single-call frequency data. The 10-dimensional single-call data were reduced to four principal components to match the explained variance (98.4%; Table S11) in the 4-call analysis. The single-call analysis showed a low false positive rate of 0.07 in stage 3, aligning closely with the theoretically expected value of 0.05 for random deviations. The false negative rate remained at 0.59, which is close to chance (21/34 = 0.617), indicating poor discriminative performance (Figure 4E). These findings underscore the developmental divergence of 4-call phrase patterns in VPA families, highlighting their potential as non-invasive biomarkers for ASD-like pups and their families. Notably, this utility is achieved without the need to identify the individual producing each call, a step that typically incurs significant costs in conventional vocal analysis, making this approach both efficient and practical.

### Parental weight loss correlates with kinship vocalisation deviation

The relationship between parental caregiver stress, indicated by weight loss, and deviations in kinship vocal communication, measured by the frequency-based divergence of 4-call phrases, was examined using a correlation analysis. In VPA families, a significant negative correlation was observed for both fathers and mothers (model #5 in Table S12; father, fix effect = −3.841, DR = 5.80, df = 1, p = 0.015; mother, fix effect = −3.093, DR = 5.05, df = 1, p = 0.024, LRT vs. model #2), indicating that greater weight loss was associated with more pronounced vocalisation deviations. No such correlation was found in UE families (model #1 in Table S12; Figure 4F). These findings suggest that stress-related body weight loss in parents is linked to atypical kinship vocal patterns in VPA families.

## Discussion

This study revealed that prenatal VPA-exposure not only affects individual marmosets but also disrupts kinship dynamics, characterised by premature locomotion independence, atypical vocalisation patterns, and significant paternal weight loss. VPA pups exhibited early independent behaviour even during the developmental phase, whereas control pups relied on parental transport. The nature of kinship vocalisations in VPA families may reflect heightened stress among family members, marked by unnatural call repetitions and minimal changes across the pre-, during-, and post-weaning periods. The frequency of call phrases in VPA families significantly deviated from the convergent patterns observed in control families after weaning. This deviation enabled the successful identification of VPA families, suggesting the potential use of kinship vocal metrics as non-invasive biomarkers. Furthermore, the inverse correlation between call phrase deviation and parental body weight highlights how altered communication may contribute to caregiver stress within the family. These findings underscore the value of prenatally VPA-exposed marmosets as a model for studying both individual ASD phenotypes and the broader impact of ASD-like traits on family dynamics, with kinship vocalisations offering a sensitive tool for assessing stress and communication impairments.

VPA pups exhibited premature locomotion independence despite having weight trajectories, a key indicator of physical development, comparable to those of the control groups. This early independence suggests disruptions in parent–pup communication during the early developmental stages. We have previously reported that VPA pups exhibit reduced social gaze and social attention by PM 3, including impairments in attachment formation,(20) and the findings of the present are consistent with these observations. Such behaviour may reflect ASD-like phenotypes, potentially mirroring hypersensitivity or hypo-responsiveness to physical contact observed in early childhood, as well as challenges in attachment formation.(30) To better understand the implications of early independence, it is essential to investigate whether this behaviour arises naturally or is influenced by the novel recording environment and whether it reflects the avoidance of parental carrying by the pup or reduced facilitation by parents. These findings emphasise the need for behavioural metrics, such as kinship vocalisations, to assess caregiver stress and family dynamics, particularly in later developmental stages.

The limited developmental changes in kinship vocalisations in VPA families were evident in their failure to exhibit typical vocal maturation observed in control families. Although the initial vocalisation patterns during the pre-weaning stage (PM 1–2.5) were comparable between the VPA and control families, significant differences emerged during later stages (PM 4–5.5). This parallels a key feature of ASD in humans, where language delays become apparent between 12 and 24 months of age as social demands increase.(31) In VPA families, critical traits such as declining call frequency, maturing call repetition patterns, and developmental changes in phrase distribution were significantly reduced, resulting in kinship vocalisations in an immature state. This developmental stagnation, combined with the paradox of premature locomotion independence, likely heightened caregiving stress and disrupted kinship dynamics. Such disruptions mirror the broader features of ASD, in which impaired communication and increased caregiving challenges significantly impact the kinship system.(32) These findings underscore the utility of VPA-exposed marmosets as a model not only for individual ASD-like traits but also for family-level dynamics, providing new insights into the interplay between communication impairments and caregiving stress in the context of ASD.

Further analysis revealed significant disruptions in the composition of calls across hierarchical levels: single calls, paired calls, and extended phrases. In the single-call analysis, trill calls accounted for half of all calls, suggesting that the recordings captured the affiliative social mood. This aligns with the findings that *phee* calls dominate in isolated conditions,(23) whereas the number of *trill* calls increases in affiliative contexts.(24) However, VPA families exhibited a pattern resembling isolated conditions, with an increased number of phee calls and a decreased number of *trill* calls, suggesting weakened social communication. The decline in the frequency of *ock* calls, often associated with contact behaviour,(22) further reflects reduced communication within the kinship system. Additionally, the occasional presence of ambiguous twitter calls in VPA families, which were almost absent in control families, underscores the broader disruption in communication dynamics caused by VPA exposure. These findings highlight the social and emotional effects of VPA exposure and provide insights into how ASD-like traits disrupt natural kinship communication.

Paired calls were analysed based on repetition and the ICI length, revealing novel insights into kinship vocal dynamics. In UE families, *trill*-*trill* pairs, which are strongly associated with affiliative interactions, were consistently predominant, replicating the findings of previous studies. By contrast, VPA families displayed deviant and less affiliative repetitions such as *trillphee*-*trillphee* and *chirp*-*chirp*. These atypical patterns may reflect tendencies analogous to echolalia or self-repetition, which are the hallmarks of ASD language characteristics.(33) Furthermore, our analysis highlighted the critical role of extended sequences consisting of four or more calls in distinguishing between VPA and control families. This distinction holds regardless of whether the sequences represent individual vocalisations or social exchanges, underscoring the need to extend the vocal structures in kinship communication analyses.(34) These findings provide a deeper understanding of the relationship between vocal communication impairments and ASD-like traits in VPA-exposed marmosets, reinforcing the utility of this model for studying both the individual- and family-level dynamics associated with ASD.

This study has several limitations. Missing data due to health conditions and experimental conflicts led to an incomplete repeated-measures design, which was addressed using linear mixed models and LRTs to account for individual and parental effects. Constraints in the soundproof room and experimental cage limited behavioural observations to basic metrics, such as whether the pups were carried or moving independently. Future research incorporating multi-camera systems and machine learning-based grammatical analyses of behaviour could provide deeper insights into kinship dynamics.(35)(36) Although our decision to avoid linking vocalisations to specific individuals preserved natural social interactions, it precluded detailed analyses of interactive versus solitary communication. Advances in machine learning enabling individual identification from vocal data offer potential solutions.(37) Nevertheless, label-free vocal analysis remains advantageous for clinical applications because it reduces labour and respects privacy.

In summary, this study demonstrates that prenatal VPA exposure disrupts both individual development and kinship dynamics in marmosets, characterised by premature locomotion independence, immature vocal communication patterns, and heightened caregiving stress. The observed stagnation in kinship vocalisation development, coupled with atypical pup locomotion, highlights the broader impact of ASD-like traits on family interactions. By leveraging non-invasive vocal metrics without individual identification, this study offers a scalable approach for assessing social dynamics, with potential applications as a biomarker for ASD traits. These findings provide valuable insights into the interplay between individual phenotypes and family systems and lay the groundwork for future research to refine ASD models and their translation into human clinical contexts.

## Materials and methods

### Subjects

All experimental and animal care procedures were approved by the Animal Research Committee of the National Center of Neurology and Psychiatry (NCNP), Tokyo, Japan, and the National Institute of Radiological Sciences, Chiba, Japan, and were performed in accordance with the United States National Institutes of Health Guide for the Care and Use of Laboratory Animals (NIH Publication Nos. 80–23) and the Guide for Care and Use of Laboratory Primates published by the National Institute of Neuroscience, NCNP.

In this study, we used 16 marmosets born from nine litters along with their parents (six males and five females; Figure 1B). The parental experience with birth and caregiving ranged from 1 to 7 times (mean ± standard deviation (SD): mother, 3.7 ± 1.9; fathers, 4.2 ± 2.0). The age of the mothers at the time of birth was 5.7 ± 1.8 years, and the age of the fathers was 7.8 ± 3.9 years (Table S1). The experimental period lasted from the birth of the pups (PM 0) to PM 5.5, during which time the parents and pups were housed together in the same cage (500 × 600 × 800 mm [width × depth × height]). Marmosets and their dams were housed in family cages and provided with food (CMS-1, CREA-Japan Inc., Tokyo, Japan) and water ad libitum. The subjects were kept at room temperature of 29 ± 2.0 °C and maintained on a 12 h:12 h light–dark cycle. The lights in the breeding room were turned on at 7:00 am and off daily at 7:00 pm. The marmosets in the facility were familiar with human contact and approached the experimenter to obtain food rewards without hesitation.

### VPA treatment

Of the nine litters, five were obtained by administering VPA to the mothers during pregnancy, resulting in VPA-exposed offspring. These parent–offspring groups were defined as the VPA family group. The remaining four litters were from the mothers that were not exposed to VPA and served as the unexposed (UE) control family group. VPA marmosets were obtained using the same procedure as previously described.13,18 Briefly, the dams were mated in their paired cages, and their blood progesterone levels were periodically monitored to determine the timing of pregnancy. Blood samples (<0.3 mL) were collected twice weekly from the femoral vein of un-anaesthetised animals placed in a restrainer (CL-4532, CLEA, Japan, Inc.). The dams received seven intragastric administrations of VPA sodium salt (200 mg/kg/day; Sigma-Aldrich, St. Louis, MO, USA) from days 60 to 66 after conception (Figure 1A). VPA was dissolved in 10% glucose solution immediately prior to administration.

### Body weight analysis

The body weights of the pups and parents were measured regularly during routine health checks. Pup weight comparisons, which had minimal missing data, were analysed using repeated-measures ANCOVA with the model weight ∼ PM + treatment + PM:treatment + error (pup) (Figure 1C). The parental body weight was calculated as a monthly average and scaled for comparison, with the period from PM −3 to −1 normalised to 1. Data from specific parent pairs were excluded from the analysis: father KU, who died suddenly during the caregiving period for pups UE2 and UE3, and the parent pairs CH and NI, who lacked experience with UE pups. Based on these criteria, body weight data were collected from the birth experiences of all the other parents (Table S2). Parental body weight group comparisons were conducted using repeated-measures ANCOVA, as was performed for pups (Figure 1E). When data sufficiency was allowed, further analysis was performed using Tukey’s HSD test for pairwise comparisons (Figure 1G).

### Vocal recording and call classification

Kinship vocalisations were recorded using a video recorder (HDR-CX630V, SONY) with the parents and one pup inside their home cage, which was moved to a soundproof room. The recordings were taken between 10:00 am and 1:00 pm. The recording began immediately after the door was closed and continued for 30 min, during which time the animals were free to move within the cage. Recordings were conducted every 1–2 weeks from the pups’ PM 1 to 5.5. However, owing to the parents’ health conditions and conflicts with other experiments, there were missing data; thus, the habituation of the pairs to the recording environment was not standardised. By using Syrinx version 2.4i (https://syrinxpc.com/) and R package seewave version 2.2.3,(38) all recorded calls were visually inspected on a spectrogram and manually classified by an expert and recorded their starting and ending timestamps, referencing characteristic features.

### Locomotion analysis

During the recording sessions, whether pups were being carried by a parent was determined through video observation. Instances where the pup’s limbs were completely off the floor or walls, relying entirely on either the father or mother for transport, were categorised as ‘carried’. All other instances were classified as ‘independent’. The classification relied solely on top-view video footage, and ambiguous situations—such as unclear carried states or when the pup moved into blind areas—were included in the ‘independent’ category. The initial independent time around PM 1 was compared between the groups using the Brunner–Munzel test to assess the median differences nonparametrically.

### LMM analysis

Owing to the irregular occurrence of missing data, applying a simple repeated-measures analysis of variance (ANOVA) was not feasible. Therefore, hypothesis testing was conducted using the ANCOVA, and model selection was performed using an LMM. In the LMM analysis, pups’ age and VPA treatment conditions were regressed against various vocalisation data, with pup sex, pup ID, and parent ID included as random effects. We ensured the robustness of the LMM analysis by conducting LRTs based on the chi-square distribution of likelihood ratios between nested models. All nested models are presented in Tables S3 and S12. The best model was selected based on the AIC to assess correlations between pups’ age and significance of the VPA treatment effect.

### Phrasing distribution analysis

For two consecutive calls, the interval between the end of the pre-call and the start of the post-call was defined as the ICI. Calls were considered related when the ICI was ≤ 30 s. Among related calls, those with an ICI ≤ 10 s were classified as ‘short ICI,’ whereas those with an ICI >10 s were classified as ‘long ICI’ (Figure 2A). Additionally, when aggregating call phrases consisting of three or more calls, only those with all ICIs ≤ 30 s were included in the analysis (Figure 3).

To quantify the change in phrase diversity between two vocal recording sessions, we adopted the JSD. JSD represents the expected difference between two frequency distributions, resolving the asymmetry present in the simple expectation of the Kullback–Leibler divergence (KLD). The JSD between the two frequency distributions *P* and *Q* is defined as

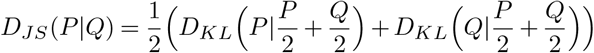

Where *D*_*KL*_ represents the KLD. The KLD from state *P* to *Q* is calculated for each phrase *a*_*i*_ as follows:

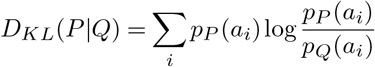

where *p*_*P*_ (*ai*) and *p*_*Q*_(*ai*) denote the probability of phrase *a*_*i*_ in distributions *P* and *Q*, respectively. The KLD quantifies the expected gain of self-information *I* when moving from distribution *P* to *Q*. The self-information *I* of state *A* = {*a*_1_, *a*_2_, …} is defined as follows:

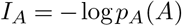

Finally, the entropy *H* of state *A* is the expected value of *I*_*A*_ given by the following equation:

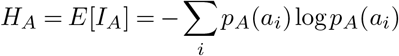

The JSD values were computed using the jsd() function in the philentropy package of R.(39)

### Discriminant analysis

In the discriminant analysis shown in Figs. 4B–F, the Mahalanobis distance was calculated using the stats::mahalanobis() function, PCA was performed using the stats::prcomp() function, and the JSD was computed using the philentropy::jsd() function. The data matrix, consisting of the number of occurrences of 4-call phrases with ICIs ≤ 30 across all PMs, was reduced to a score matrix of the first to fifth principal components using PCA based on the covariance matrix, which accounted for 98.4% of the data variance (Table S8). PCA was also performed on the frequency of single calls for comparison; in this case, the first to fourth principal components were used to match the data variance (98.4%, Table S10). These data were then divided into three stages—stage 1 (PM 1–2.5), stage 2 (PM 2.5–4), and stage 3 (PM 4–5.5)—and Mahalanobis distances were calculated for each stage with the UE group as the reference. The threshold for identifying outliers was set to 11.0705 for 4-call phrases and 9.4877 for single calls, corresponding to the 95th percentile of the chi-square distribution with five and four degrees of freedom, respectively.

The Mahalanobis distance of data point A from group B is defined as follows:

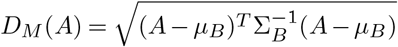

where *µ*_*B*_ is the mean vector of group *B*, and 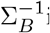 is the inverse of the covariance matrix of group *B*. The false negative and false positive ratios were defined as the proportions of VPA and UE family data points, respectively, within a specified period where the Mahalanobis distance did not exceed (false negative) or exceeded (false positive) the threshold, which was determined as the 95% distribution range expected from UE family data points, indicating the absence or presence of observed deviations.

### Statistical Analysis

All data processing and statistical analyses were performed using R version 4.3.3. For data processing, we employed {tidyverse} 2.0.0(40) and {data.table} 1.15.0. Data visualisation was performed using {ggplot2} 3.5.1. Statistical analyses were performed using {rstatix} for type II ANOVA (data ∼ treatment * individuals), ANCOVA (data ∼ treatment * PM), and F-test (data ∼ PM) for regression analysis, {lawstat} 3.6 for non-parametric hypothesis testing, {lme4} 1.1–35.5, for LMM analysis, and a custom function for LRT based on the LMM results.

## ACKNOWLEDGEMENTS

We thank R. Saito, S. Okamura, A. Tsuchiya, I. Yamamoto, and Y. Nishimura for technical assistance. We also thank Dr. J. Noguchi (NCNP), Dr. K. Yoshitake (NCNP), Dr. K. Shimatani (Institute of Statistical Mathematics), Dr. S. Nakamura (Tokyo University of Agriculture and Technology), and Dr. T. Minamimoto (National Institutes for Quantum Science and Technology) for their comments on an earlier version of the manuscript.

This study was supported by JSPS Research Fellowships for Young Scientists 14J10961 (to KM), MEXT/JSPS KAKENHI under the grant Number J22K07338 (to KM), AMED under grant numbers JP24wm0625124 (to KM), JP24wm0625206 (to HM), 22dm027066h004 (to NI), and the Intramural Research Grant for Neurological and Psychiatric Disorders from NCNP under grant numbers 2-7 and 5-8 (to NI).

