## Supplemental data for "Altered kinship vocal dynamics in marmosets with valproic acid–induced model of autism"

**Table S1. Pups and parents with recorded kinship vocalisations**

| Pup | Sex | Birth | Mother | Age (y) | NOP | Father | Age (y) | NOP |
|---|---|---|---|---|---|---|---|---|
| UE1 | m | 2015/01/15 | CH | 4.5 | 3 | KU | 8.1 | 5 |
| UE2 | f | 2015/07/13 | CH | 5.0 | 4 | KU | 8.6 | 6 |
| UE3 | m | 2015/07/13 | CH | 5.0 | 4 | KU | 8.6 | 6 |
| UE4 | m | 2015/06/17 | SA | 9.3 | 7 | IW | 9.7 | 7 |
| UE5 | m | 2015/06/17 | SA | 9.3 | 7 | IW | 9.7 | 7 |
| UE6 | m | 2015/08/16 | BE | 5.0 | 1 | DO | 3.3 | 1 |
| UE7 | f | 2015/08/16 | BE | 5.0 | 1 | DO | 3.3 | 1 |
| VPA1 | m | 2016/04/21 | CH | 5.8 | 5 | NI | 10.8 | 5 |
| VPA2 | m | 2016/04/21 | CH | 5.8 | 5 | NI | 10.8 | 5 |
| VPA3 | f | 2014/12/22 | SA | 8.8 | 6 | IW | 9.2 | 6 |
| VPA4 | m | 2016/02/13 | BE | 5.5 | 2 | DO | 3.8 | 2 |
| VPA5 | m | 2016/02/13 | BE | 5.5 | 2 | DO | 3.8 | 2 |
| VPA6 | m | 2015/06/04 | AN | 5.3 | 3 | SH | 3.7 | 3 |
| VPA7 | f | 2015/06/04 | AN | 5.3 | 3 | SH | 3.7 | 3 |
| VPA8 | m | 2015/04/30 | BU | 3.6 | 3 | CO | 14.2 | 4 |
| VPA9 | m | 2015/04/30 | BU | 3.6 | 3 | CO | 14.2 | 4 |

NOP, number of parenting experiences

**Table S2. Parenting experience of the parents used in the body weight analysis**

| Mother | NOP | Father | NOP | Child birth | Treatment |
|---|---|---|---|---|---|
| BU | 1 | CO | 2 | 2014/4/9 | UE |
| BU | 2 | CO | 3 | 2014/10/11 | UE |
| BU | 3 | CO | 4 | 2015/4/30 | **VPA** |
| BE | 1 | DO | 1 | 2015/8/16 | UE |
| BE | 2 | DO | 2 | 2016/2/13 | **VPA** |
| BE | 3 | DO | 3 | 2016/8/12 | UE |
| BE | 4 | DO | 4 | 2017/3/19 | **VPA** |
| BE | 5 | DO | 5 | 2019/6/24 | **VPA** |
| SA | 1 | IW | 1 | 2011/9/29 | UE |
| SA | 2 | IW | 2 | 2012/2/29 | UE |
| SA | 3 | IW | 3 | 2012/10/20 | **VPA** |
| SA | 4 | IW | 4 | 2013/4/18 | UE |
| SA | 5 | IW | 5 | 2014/6/27 | UE |
| SA | 6 | IW | 6 | 2014/12/22 | **VPA** |
| SA | 7 | IW | 7 | 2015/6/17 | UE |
| SA | 8 | IW | 8 | 2016/1/9 | **VPA** |
| SA | 9 | IW | 9 | 2016/7/30 | UE |
| SA | 10 | IW | 10 | 2017/1/25 | UE |
| SA | 11 | IW | 11 | 2017/8/19 | UE |
| CH | 1 | KU | 3 | 2013/11/21 | **VPA** |
| CH | 2 | KU | 4 | 2014/6/23 | **VPA** |
| CH | 3 | KU | 5 | 2015/1/15 | UE |
| CH | 4 | KU | 6 | 2015/7/13 | UE |
| AN | 1 | SH | 1 | 2014/1/4 | **VPA** |
| AN | 2 | SH | 2 | 2014/6/7 | **VPA** |
| AN | 3 | SH | 3 | 2015/6/4 | **VPA** |
| AN | 4 | SH | 4 | 2016/7/25 | UE |
| AN | 5 | SH | 5 | 2017/1/25 | **VPA** |

NOP, number of parenting experiences

**Table S3. Nested models in the linear mixed model analysis**

| Model | Formula |
|---|---|
| **#1** | Call ~ 1 |
| **#2** | Call ~ 1 + (PM \| treatment) |
| **#3** | Call ~ 1 + (PM \| sex) |
| **#4** | Call ~ 1 + (PM \| pup) |
| **#5** | Call ~ 1 + (PM \| parents) |
| **#6** | Call ~ treat |
| **#7** | Call ~ treat + (treatment \| sex) |
| **#8** | Call ~ treat + (treatment \| pup) |
| **#9** | Call ~ treat + (treatment \| parents) |
| **#10** | Call ~ PM |
| **#11** | Call ~ PM + (PM \| treat) |
| **#12** | Call ~ PM + (PM \| sex) |
| **#13** | Call ~ PM + (PM \| pup) |
| **#14** | Call ~ PM + (PM \| parents) |
| **#15** | Call ~ treat + PM + treatment:PM |
| **#16** | Call ~ treat + PM + treatment:PM + (PM \| sex) |
| **#17** | Call ~ treat + PM + treatment:PM + (PM \| pup) |
| **#18** | Call ~ treat + PM + treatment:PM + (PM \| parents) |
| **#19** | Call ~ treat + PM + treatment:PM + (treatment \| sex) |
| **#20** | Call ~ treat + PM + treatment:PM + (treatment \| pup) |
| **#21** | Call ~ treat + PM + treatment:PM + (treatment \| parents)) |

The variables on the right side of the '|' symbol indicate the random effects introduced by the factors on the left side. The ':' symbol represents interactions between the preceding and following variables. 'Call' is a parameter related to calls,

including the call frequency and population ratio; PM indicates the age of the pup‘ in postnatal months; and treatment indicates the VPA exposure condition of the pup, either VPA-exposed or unexposed.
VPA, valproic acid; PM, postnatal months.

**Table S4. AIC of the LMM analysis in the correlation between call frequency and PM**

| Treatment | Model | AIC | dAIC |
|---|---|---|---|
| UE | #1 | 666.40 | 31.86 |
| | #3 | 659.12 | 24.58 |
| | #4 | 645.27 | 10.72 |
| | #5 | 655.07 | 20.52 |
| | #10 | 661.33 | 26.78 |
| | #12 | 649.08 | 14.54 |
| | **#13** | **634.55** | **0.00** |
| | #14 | 644.81 | 10.27 |
| VPA | #1 | 863.83 | 26.20 |
| | #3 | 861.21 | 23.58 |
| | #4 | 847.65 | 10.02 |
| | #5 | 844.94 | 7.30 |
| | #10 | 865.83 | 28.19 |
| | #12 | 854.94 | 17.31 |
| | #13 | 841.04 | 3.41 |
| | **#14** | **837.63** | **0.00** |

VPA, valproic acid; UE, unexposed; LMM, linear mixed model; AIC, Akaike information criterion.

**Table S5. Definitions of the nice main call types in marmoset kinship vocalisations**

| Call type | |
|---|---|
| ock | The frequency is generally below 4 kHz, with a peak-like shape. The sound resembles a clicking or coughing noise. |
| tsik | A brief call, initially exceeding 12 kHz in frequency, followed by a line that extends from low to high frequencies on the spectrogram towards the end. |
| egg | Similar to ‘ock’ call but composed of intermittent harmonics. It is sometimes vocalised in a series. |
| phee | A strong, prolonged call lasting more than 1 s, sometimes extending beyond 2 s. The frequency is typically around 8 kHz, although there is some variation. |
| peep | A short, weak call with minimal frequency variation between the start and the end or a slight upward shift of less than 2 kHz. It is sometimes emitted in succession but may also occur as a single call. |
| trill | A distinctive call accompanied by waveform oscillations, showing a wide range of variation in duration, intensity, and pitch. |
| trillphee | A single call formed by a trill immediately followed by a phee-like phrase, connected seamlessly. |
| chirp | A short call where the ending frequency is approximately 2 kHz lower than the starting frequency. It is occasionally observed in succession. |
| twitter | A highly distinctive call composed of a series of very short sounds occurring in quick succession. |

**Table S6. List of dAICs in the LMM analysis of each call type**

| Model | chirp | egg | ock | other | peep | phee | trill | trillphee | tsik | twitter |
|---|---|---|---|---|---|---|---|---|---|---|
| #1 | 28.6 | 1.7 | 15.6 | 10.3 | 74.2 | 23.3 | 59.4 | 55.3 | 13.7 | 0.7 |
| #2 | 20.5 | 3.7 | 16.2 | 10.7 | 70.9 | 22.3 | 46.5 | 48.5 | 13.3 | 6.4 |
| #3 | 23.4 | 3.7 | 16.7 | 10.9 | 73.2 | 25.3 | 53.1 | 50.0 | 13.7 | 6.4 |
| #4 | 11.9 | 3.4 | 4.2 | 9.7 | 27.7 | 10.7 | 20.3 | 14.6 | 10.9 | **0** |
| #5 | 16.1 | 3.4 | 3.0 | 10.6 | 12.7 | 17.1 | 39.5 | 17.2 | 9.7 | 4.8 |
| #6 | 29.8 | 3.5 | 12.2 | 10.4 | 72.6 | 19.9 | 55.6 | 56.6 | 14.1 | 2.3 |
| #7 | 26.2 | 0.2 | 11.2 | 4.3 | 63.2 | 18.8 | 45.2 | 53.8 | 6.7 | 4.8 |
| #8 | 21.1 | 0.0 | 3.0 | 1.8 | 36.6 | 8.8 | 17.8 | 21.8 | 4.8 | 4.7 |
| #9 | 25.3 | **0** | 0.7 | 4.1 | 38.5 | 6.8 | 39.3 | 22.9 | 4.9 | 4.8 |
| #10 | 19.0 | 3.3 | 16.0 | 12.2 | 75.3 | 23.7 | 52.7 | 45.1 | 14.7 | 1.6 |
| #11 | 18.1 | 5.0 | 17.6 | 10.5 | 68.5 | 22.9 | 41.2 | 46.1 | 12.1 | 10.2 |
| #12 | 20.5 | 5.0 | 18.8 | 10.8 | 70.7 | 26.7 | 47.4 | 46.9 | 12.3 | 10.2 |
| #13 | 6.4 | 4.8 | 4.9 | 9.5 | 25.0 | 11.1 | 13.9 | 13.3 | 9.9 | 4.0 |
| #14 | 11.5 | 4.6 | 3.7 | 10.6 | 8.6 | 17.5 | 33.5 | 16.6 | 8.1 | 8.8 |
| #15 | 17.8 | 7.2 | 13.8 | 14.1 | 74.4 | 19.8 | 47 | 45.4 | 16.7 | 3.5 |
| #16 | 11.6 | 1.4 | 13.6 | 2.4 | 56.4 | 18.7 | 26.1 | 41.3 | 3.2 | 8.7 |
| #17 | **0** | 1.1 | 0.1 | 1.7 | 13.7 | 6.4 | **0** | 9.0 | 1.4 | 2.4 |
| #18 | 5.0 | 1.0 | **0** | 2.4 | **0** | **0** | 22.8 | **0** | **0** | 7.1 |
| #19 | 11.2 | 1.4 | 13.0 | 2.4 | 56.4 | 18.7 | 26.6 | 41.3 | 3.2 | 8.7 |
| #20 | 8.7 | 1.3 | 4.3 | **0** | 29.8 | 8.9 | 4.6 | 18.0 | 1.7 | 8.6 |
| #21 | 11.5 | 1.3 | 1.8 | 2.3 | 30.3 | 7.5 | 21.9 | 14.8 | 1.6 | 8.7 |

dAIC, delta Akaike information criterion; LMM, linear mixed model

### Table S7. LMM analysis results for repeated and non-repeated calls

| ICI | Bigram type | Treatment | Model | AIC | dAIC |
|---|---|---|---|---|---|
| Short | Repeated | UE | #1 | -37.33 | 12.05 |
| | | | #3 | -38.93 | 10.44 |
| | | | #4 | -48.51 | 0.86 |
| | | | #5 | -47.82 | 1.55 |
| | | | **#10** | -49.38 | **0** |
| | | | #12 | -31.67 | 17.70 |
| | | | #13 | -42.93 | 6.44 |
| | | | #14 | -40.95 | 8.42 |
| Long | Repeated | VPA | #1 | -35.73 | 15.94 |
| | | | #3 | -23.99 | 27.68 |
| | | | #4 | -42.89 | 8.78 |
| | | | **#5** | -51.68 | **0** |
| | | | #10 | -33.75 | 17.92 |
| | | | #12 | -12.97 | 38.70 |
| | | | #13 | -32.29 | 19.38 |
| | | | #14 | -41.28 | 10.39 |
| Short | Non-repeated | UE | #1 | -27.86 | 7.19 |
| | | | #3 | -21.21 | 13.84 |
| | | | #4 | -21.38 | 13.66 |
| | | | #5 | -22.26 | 12.79 |
| | | | **#10** | -35.05 | **0** |
| | | | #12 | -14.22 | 20.83 |
| | | | #13 | -16.01 | 19.04 |
| | | | #14 | -15.41 | 19.63 |
| Long | Non-repeated | VPA | **#1** | -91.43 | **0** |
| | | | #3 | -79.18 | 12.24 |
| | | | #4 | -83.55 | 7.88 |
| | | | #5 | -87.90 | 3.52 |
| | | | #10 | -89.84 | 1.58 |
| | | | #12 | -70.27 | 21.158 |
| | | | #13 | -71.95 | 19.47 |
| | | | #14 | -76.58 | 14.84 |

AIC, Akaike information criterion; dAIC, delta AIC; LMM, linear mixed model; ICI, VPA, valproic acid; UE, unexposed.

**Table S8. List of dAICs in the LMM analysis of each repeated call with a short ICI**

| Model | ock-ock | tsik-tsik | egg-egg | phee-phee | peep-peep | trill-trill | trillphee-trillphee | chirp-chirp | twitter-twitter | other-other |
|---|---|---|---|---|---|---|---|---|---|---|
| 1 | 1.98 | NA | **0** | **0** | 50.2 | 28.7 | 3.1 | 0.6 | NA | 6.0 |
| 2 | 12.9 | NA | 12.8 | 12.7 | 49.9 | 23.2 | 14.4 | 11.1 | NA | 15.1 |
| 3 | 14.3 | NA | 12.9 | 12.8 | 58.3 | 40.0 | 17.3 | 12.1 | NA | 13.8 |
| 4 | 10.6 | NA | 10.7 | 12.8 | 4.6 | 15.5 | 15.9 | 4.2 | NA | 4.2 |
| 5 | 12.4 | NA | 11.9 | 12.8 | **0** | 7.4 | 15.5 | 2.7 | NA | 7.0 |
| 6 | 3.64 | NA | 1.2 | 0.8 | 37.6 | 9.6 | 0.2 | **0** | NA | 5.3 |
| 7 | 21.5 | NA | 19.0 | 18.5 | 45.5 | 22.9 | 21.2 | 15.5 | NA | 22.1 |
| 8 | 21.5 | NA | 19.6 | 18.3 | 9.4 | 8.3 | 22.1 | 10.8 | NA | 17.8 |
| 9 | 20.9 | NA | 19.6 | 18.7 | 3.2 | **0** | 22.2 | 6.6 | NA | 21.7 |
| 10 | **0** | NA | 2.0 | 1.1 | 47.0 | 30.1 | 2.8 | 2.2 | NA | **0** |
| 11 | 19.3 | NA | 22.2 | 21.3 | 55.6 | 31.0 | 22.6 | 18.9 | NA | 19.6 |
| 12 | 20 | NA | 21.2 | 21.3 | 62.6 | 47.2 | 25.8 | 19.8 | NA | 19.6 |
| 13 | 16.7 | NA | 19.9 | 21.5 | 8.2 | 22.8 | 24.6 | 11.9 | NA | 10.3 |
| 14 | 17.9 | NA | 21.1 | 21.3 | 4.1 | 12.7 | 24.3 | 9.7 | NA | 12.9 |
| 15 | 1.14 | NA | 3.4 | 3.3 | 38.6 | 13.3 | **0** | 3.0 | NA | 3.87 |
| 16 | 32.4 | NA | 34.6 | 34.6 | 61.9 | 39.3 | 36.9 | 29.4 | NA | 32.6 |
| 17 | 28.1 | NA | 34.5 | 34.4 | 14.0 | 25.7 | 38.1 | 22.1 | NA | 23.9 |
| 18 | 29.8 | NA | 35.4 | 34.5 | 15.3 | 22.2 | 38.1 | 19.1 | NA | 24.8 |
| 19 | 32.2 | NA | 34.5 | 34.4 | 57.3 | 37.2 | 37.1 | 29.4 | NA | 32.6 |
| 20 | 32.2 | NA | 35.9 | 34.4 | 19.6 | 22.4 | 38.0 | 23.8 | NA | 24.6 |
| 21 | 30.4 | NA | 35.9 | 34.6 | 12.9 | 13.9 | 38.2 | 19.6 | NA | 24.8 |

The data for tsik-tsik and twitter-twitter were insufficient for the LMM analysis. dAIC, delta AIC; LMM, linear mixed model.

**Table S9. Counts of representative 4-call phrases at each developmental stage**

| Lank | Stage | PM | phrase | UE | VPA | total |
|---|---|---|---|---|---|---|
| 1 | 1 | 1-2.5 | trill-trill-trill-trill | 2128 | 693 | 2821 |
| 2 | 1 | 1-2.5 | chirp-chirp-chirp-chirp | 200 | 227 | 427 |
| 3 | 1 | 1-2.5 | trill-trill-trillphee-trill | 173 | 84 | 257 |
| 4 | 1 | 1-2.5 | trill-trillphee-trill-trill | 169 | 80 | 249 |
| 5 | 1 | 1-2.5 | trillphee-trill-trill-trill | 172 | 76 | 248 |
| 6 | 1 | 1-2.5 | peep-peep-peep-peep | 72 | 173 | 245 |
| 7 | 1 | 1-2.5 | peep-trill-trill-trill | 147 | 84 | 231 |
| 8 | 1 | 1-2.5 | trill-trill-trill-trillphee | 161 | 65 | 226 |
| 9 | 1 | 1-2.5 | trill-trill-trill-peep | 132 | 73 | 205 |
| 10 | 1 | 1-2.5 | trill-trill-peep-trill | 136 | 68 | 204 |
| 1 | 2 | 2.5-4 | trill-trill-trill-trill | 807 | 629 | 1436 |
| 2 | 2 | 2.5-4 | peep-peep-peep-peep | 39 | 996 | 1035 |
| 3 | 2 | 2.5-4 | chirp-chirp-chirp-chirp | 9 | 150 | 159 |
| 4 | 2 | 2.5-4 | peep-trill-trill-trill | 75 | 73 | 148 |
| 5 | 2 | 2.5-4 | trill-trill-trill-peep | 68 | 72 | 140 |
| 6 | 2 | 2.5-4 | trill-peep-trill-trill | 62 | 65 | 127 |
| 7 | 2 | 2.5-4 | trill-trill-peep-trill | 62 | 64 | 126 |
| 8 | 2 | 2.5-4 | chirp-trill-trill-trill | 55 | 50 | 105 |
| 9 | 2 | 2.5-4 | trill-chirp-trill-trill | 51 | 45 | 96 |
| 10 | 2 | 2.5-4 | trill-trill-chirp-trill | 55 | 39 | 94 |
| 1 | 3 | 4-5.5 | peep-peep-peep-peep | 2 | 1800 | 1802 |
| 2 | 3 | 4-5.5 | trill-trill-trill-trill | 283 | 461 | 744 |
| 3 | 3 | 4-5.5 | chirp-chirp-chirp-chirp | 19 | 109 | 128 |
| 4 | 3 | 4-5.5 | tsik-tsik-tsik-tsik | 0 | 120 | 120 |
| 5 | 3 | 4-5.5 | egg-tsik-egg-tsik | 0 | 72 | 72 |
| 6 | 3 | 4-5.5 | trillphee-trill-trill-trill | 19 | 51 | 70 |
| 7 | 3 | 4-5.5 | trill-trillphee-trill-trill | 18 | 50 | 68 |
| 8 | 3 | 4-5.5 | peep-trill-trill-trill | 22 | 44 | 66 |
| 9 | 3 | 4-5.5 | trill-trill-trillphee-trill | 16 | 49 | 65 |
| 10 | 3 | 4-5.5 | trill-trill-trill-peep | 20 | 43 | 63 |

VPA, valproic acid; UE, unexposed.

**Table S10. Summary of the principal component (PC) for the frequency of 4-call phrases**

| | **PC1** | **PC2** | **PC3** | **PC4** | **PC5** | PC6 | PC7 |
|---|---|---|---|---|---|---|---|
| Standard deviation | **124.396** | **89.248** | **13.050** | **12.686** | **11.782** | 8.553 | 7.372 |
| Variance explained | **0.637** | **0.328** | **0.007** | **0.006** | **0.005** | 0.003 | 0.002 |
| Cumulative proportion | **0.637** | **0.965** | **0.972** | **0.978** | **0.984** | 0.987 | 0.989 |

**Table S11. Summary of the principal component (PC) for the frequency of single calls**

| | **PC1** | **PC2** | **PC3** | **PC4** | PC5 | PC6 | PC7 |
|---|---|---|---|---|---|---|---|
| Standard deviation | **170.125** | **135.931** | **32.365** | **25.105** | 18.949 | 13.095 | 11.719 |
| Variance explained | **0.580** | **0.370** | **0.021** | **0.012** | 0.007 | 0.003 | 0.002 |
| Cumulative proportion | **0.580** | **0.950** | **0.971** | **0.984** | 0.991 | 0.994 | 0.997 |

**Table S12. LMM analysis results of the body weight and phrase frequency deviation rate**

| | VPA | | | | UE | | | |
|---|---|---|---|---|---|---|---|---|
| | Father | | Mother | | Father | | Mother | |
| Model | AIC | dAIC | AIC | dAIC | AIC | dAIC | AIC | dAIC |
| #1 Dev ~ 1 | 120 | 21.0 | 151 | 4.2 | -1 | **0** | -1 | **0** |
| #2 Dev ~ 1 + (BW \| parents) | 103 | 3.8 | 150 | 3.1 | 9.3 | 11 | 9.3 | 11 |
| #3 Dev ~ 1 + (BW \| pup) | 127 | 28.0 | 154 | 7.2 | 8.4 | 9.7 | 8.4 | 9.7 |
| #4 Dev ~ BW | 121 | 21.0 | 153 | 6.1 | 0.4 | 1.7 | 0.6 | 1.9 |
| #5 Dev ~ BW + (BW \| parents) | 100 | **0** | 147 | **0** | 9.5 | 11 | 9.6 | 11 |
| #6 Dev ~ BW + (BW \| pup) | 108 | 8.3 | 152 | 5.2 | 8.0 | 9.3 | 8.9 | 10 |

The variables on the right side of the '|' symbol indicate the random effects introduced by the factors on the left side. The ':' symbol represents interactions between the preceding and following variables. Dev indicates the deviation index defined as the log10 of the Mahalanobis distance from UE families; BW indicates scaled body weight.

LMM, linear mixed model; AIC, Akaike information criterion; dAIC, delta AIC; VPA, valproic acid; UE, unexposed.
